# Quantitative Relations in Protein and RNA Folding Deduced from Quantum Theory

**DOI:** 10.1101/021782

**Authors:** Liaofu Luo, Jun Lv

**Author notes:** revised version of preprint* “Folding rate of protein and RNA studied from quantum folding theory” *posted at* http://dx.doi.org/10.1101/021782.

## Abstract

Quantitative relations in protein and RNA folding are deduced from the quantum folding theory of macromolecules. It includes: deduction of the law on the temperature-dependence of folding rate and its tests on protein dataset; study on the chain-length dependence of the folding rate for a large class of biomolecules; deduction of the statistical relation of folding free energy versus chain-length; and deduction of the statistical relation between folding rate and chain length and its test on protein and RNA dataset. In the above quantum approach the influence of the solvent environment factor on folding rate has been taken into account automatically. The successes of the deduction of these new relations from the first principle and their successful comparison with experimental data afford strong evidence on the possible existence of a common quantum mechanism in the conformational change of biomolecules.

## 1 Introduction

There are huge numbers of variables in a biological system. What are the fundamental variables in the life processes at the molecular level? Since the classical works of B. Pullman and A. Pullman on nucleic acids [1], it is generally accepted that the mobile *π* electrons play an important role in the biological activities of macromolecules. However, the quantum biochemistry cannot treat a large class problems relating to the conformational variation of biological macromolecules such as protein folding, RNA folding, signal transduction and gene expression regulation, etc. In fact, for a macromolecule consisting of *n* atoms there are 3*n* coordinates if each atom is looked as a point. Apart from 6 translational and rotational degrees of freedom there are 3*n*-6 coordinates describing molecular shape. It has been proved that the bond lengths, bond angles and torsion (dihedral) angles form a complete set to describe the molecular shape. The molecular shape is the main variables responsible for conformational change. However, to our knowledge, there is no successful approach to the quantum motion of molecular shape.

For a complex system consisting of many dynamical variables the separation of slow/fast variables is the first key step in the investigation. In his synergetics Haken proposed that the long-living systems slave the short-living ones, or briefly, the slow variables slave the fast ones. He indicated that the fast variables can be adiabatically eliminated in classical statistical mechanics [2]. However, what is the slow variable for a molecular biological system? The typical chemical bond energy is several electron volts (for example, 3.80 ev for C-H bond, 3.03 ev for C-N bond, 6.30 ev for C=O dissociation). The CG hydrogen bond energy is 0.2 ev and the TA hydrogen bond energy is 0.05 ev in nucleic acids. The energy related to the variation of bond length and bond angle is in the range of 0.4-0.03 ev. While the torsion vibration energy is 0.03-0.003 ev, the lowest in all forms of biological energies. In terms of frequency, the stretching and bending frequency is 10^14^–10^13^ Hz while that for torsion is 7.5×10^12^–7.5×10^11^ Hz. Interestingly, the torsion energy is even lower than the average thermal energy per atom at room temperature (0.04 ev in 25 C); the torsion angles are easily changed even at physiological temperature. Therefore, the torsion motion can be looked as the slow variable and others including mobile *π* electron, chemical binding, stretching and bending etc are fast variables. Moreover, different from stretching and bending the torsion potential generally has several minima that correspond to several stable conformations. Therefore, it is reasonable to assume that the torsions are slaving slow variables for the biomolecule system, the molecular conformations can be defined by torsion states and the conformational change is essentially a quantum transition between them [3,4].

Although the knowledge of protein structure is in a phase of remarkably growth the dynamical mechanism of protein folding is still unclear. The molecular dynamics (MD) provides a computer method to simulate the mechanism only at the coarse grained level since it is mainly based on the classical mechanics. Simultaneously, due to the large computational cost the limited results currently obtained by atomic-level MD simulations (such as on massively parallel supercomputer Anton [5]) are still inadequate for giving an answer on the basic folding mechanism. A noted problem is why the protein folding rate always exhibits the curious non-Arrhenius temperature dependence (the logarithm folding rate is not a decreasing linear function of 1/*T*)[6]. The non-Arrhenius peculiarity was conventionally interpreted by the nonlinear temperature dependence of the configurational diffusion constant on rough energy landscapes [7], by the temperature dependence of hydrophobic interaction [8,9] or by introducing the number of denatured conformation depending on temperature to interpret the difference between folding and unfolding [6]. All these explanations are inconclusive. Recent experimental data indicated very different and unusual temperature dependencies of the folding rates existing in the system of *λ*_6-85_ mutants [10] and in some de novo designed ultrafast folding protein [11]. These unusual Arrhenius plots of ultrafast folders provide an additional kinetic signature for protein folding, indicating the necessity of searching for a new folding mechanism. Another problem is related to the Levinthal’s paradox. In a recent work Garbuzynskiy et al reported that the measured protein folding rates fall within a narrow triangle (called Golden triangle)[12]. Why protein folding rates are confined in such a narrow kinetic region? To explain these longstanding problems it seems that a novel physical model on the folding mechanism is required. Based on the idea that the molecular conformation is defined by torsion state and the folding/unfolding is essentially a quantum transition between them, through adiabatically elimination of fast variables we are able to obtain a set of fundamental equations to describe the rate of conformational transition of macromolecule [3,4]. Our experience shows that the adiabatically elimination is an effective tool to deal with the multi-scale problems such as protein folding. By use of these equations we have successfully explained the non-Arrhenius temperature dependence of the folding rate for each protein. Moreover, the statistical investigation of 65 two-state protein folding rates (which were studied by Garbuzynskiy et al) shows these fundamental equations are consistent with experimental data [13][14].

Apart from protein folding, conformation transition occurs in many other molecular biological processes. Both RNA and protein are biological macromolecules. Common themes of RNA and protein folding were indicated [15]. It is expected that both obey the same dynamical laws and have a unifying folding mechanism. In a recent work Hyeon et al reported that RNA folding rates are determined by chain length [16]. Then, a more general view on the self-assembly of proteins and RNA and some universal relations were proposed [17]. However, Hyeon’s relation of RNA folding rates vs chain length is obtained empirically and its generalization to protein is problematic. Based on quantum folding theory [4] we can make comparative studies on protein and RNA folding and deduce a theoretical formula on RNA folding rate. Our results show that the quantum theory serves a logic foundation for understanding this universality between protein and RNA folding. Moreover, the conformational quantum transition of biological molecules may provide a clue to searching for some expected quantitative unifying theory in living systems [18].

In the article we shall sketch the deduction of the general rate equation from quantum transition theory at first (in Method section). Then, main results in developing quantum folding theory to the protein and RNA folding problems will be given. It includes: deduction of the law on the temperature-dependence of folding rate and tests of the law on protein dataset; deduction of the relation of folding and unfolding parameters; study on the chain-length dependence of the fast-variable factor of the folding rate; deduction of the statistical relation of folding free energy versus chain-length; and deduction of the statistical relation between folding rate and chain length and test of the relation on protein and RNA dataset.

## 2 Materials and Methods

### 2.1 Datasets

Recently Garbuzynskiy and coworkers collected folding rate data for 69 two-state proteins [12]. Of the 69 proteins, the folding rates of 65 two-state proteins are obtained at around 25 °C. They constitute a dataset used by us to compare the theoretical vs. experimental results [13]. Hyeon and Thirumalai collected the folding rates of 27 RNA molecules [16]. They constitute the second dataset we shall use. In addition, the temperature dependence data of the folding rate for 16 proteins are collected in Table 2 of ref [13].

### 2.2 Theoretical model

Suppose the quantum state of a macromolecule is described by a wave function *M*(*θ, x*), where {*θ*} the torsion angles of the molecule and {*x*} the set of fast variables including the stretching-bending coordinates and the frontier electrons of the molecule, etc. The wave function *M*(*θ,x*) satisfies

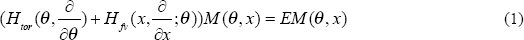

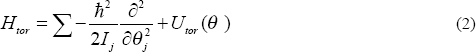

where *I_j_* denotes the inertial moment of the *j*-th torsion and the torsion potential *U_tor_* is a function of a set of torsion angles *θ*={*θ_j_*}. Its form is dependent of solvent environment of the molecule.

As is well known, the influence of solvent is a serious problem in molecular dynamics approach to any real biological problem. Suppose the interaction between water (or other solvent, ion and denaturant) molecules (their coordinates denoted by *r*) and macromolecule is *V(r,θ,x)*. Its average over *r* in a given set of experimental conditions (including chemical denaturants, solvent conditions etc.) can be expressed by 〈*V(r,θ,x))〉_av_*=*V*_1_(*θ*) + *V*_2_(*x,q*) where *V*_1_ (*θ*)is *x*-independent part of the average interaction. Then we define *U_tor_(θ)*=*U_tor,vac_(θ)* + *V*_1_(*θ*) and 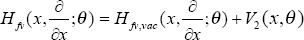 with *U_tor,vac_* the torsion potential in vacuum and *H_fv,vac_* the fast-variable Hamiltonian in vacuum. By inserting them into (1) and (2) the basic equations for the macromolecule are obtained. Thus, in the present theory the influence of the solvent environment factors has been taken into account automatically by introducing torsion potential *U_tor_* and fast-variable Hamiltonian *H_fv_* in the basic equations.

Because the fast variables change more quickly than the variation of torsion angles, the adiabatic approximation can be used. In adiabatic approximation the wave function is expressed as

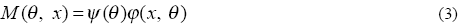

and these two factors satisfy

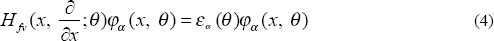

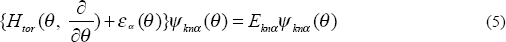

where *α* denotes the quantum number of fast-variable wave function *φ*, and (*k, n*) refer to the conformational (indicating which minimum the wave function is localized around) and the vibrational state of torsion wave function *ψ*, respectively.

Because *M (θ,x)* is not a rigorous eigenstate of Hamiltonian *H_tor_* + *H_fv_*, there exists a transition between adiabatic states that results from the off-diagonal elements

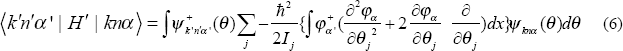

Here *H*′ is a Hamiltonian describing conformational transition. The nonadiabatic matrix element (6) can be calculated under the perturbation approximation. Through tedious calculation we obtain the rate of conformational transition [4]

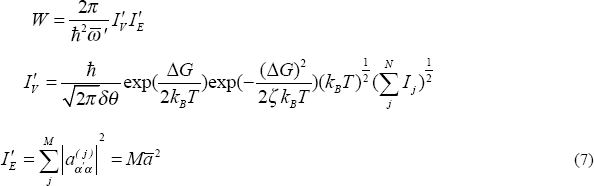

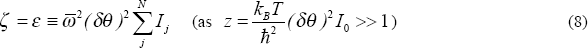

where *W* means the rate of conformational transition at given temperature *T* and solvent condition, 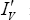 is slow-variable factor and 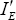 fast-variable factor, *N* is the number of torsion modes participating in a quantum transition coherently, *I_j_* denotes the inertial moment of the atomic group of the *j*-th torsion mode (*I*_0_ denotes its average hereafter), 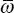 and 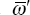 are the initial and final frequency parameters *w_j_* and 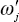 of torsion potential averaged over *N* torsion modes, respectively, *δθ* is the averaged angular shift between initial and final torsion potential. *ζ* in Eq (7) is a function of 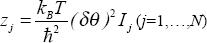 and torsion potential parameters, and is defined through a complicated combination of Bessel functions. As z>>1 *ζ* can be expressed as Eq (8), namely *ζ=ε*. The condition z>>1 is generally satisfied for most biomolecules in the following discussion. Δ*G* is the free energy decrease per molecule between initial and final states,

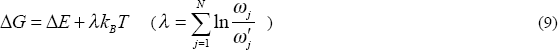

where 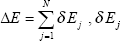the energy gap between initial and final states for the *j*-th mode, *M* is the number of torsion angles correlated to fast variables, 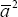 is the square of the matrix element of the fast-variable Hamiltonian operator, or, more accurately, its change with torsion angle, averaged over *M* modes,

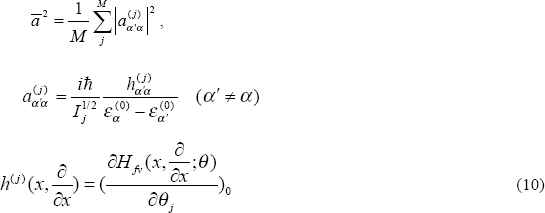

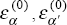 are the eigenvalues of 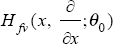

Above equations are deduced under the adiabatic approximation. The decoherence effect due to the quantum entanglement with environment can be easier to estimate because of the multiplicational form of the wave function (Eq (3)). By calculating decoherence time it was proved that the decoherence effect on molecular torsion is in the mid of electron and atom [4]. So, even if the quantum coherence for the macromolecule as a single particle may have been destroyed the coherence in the torsional degree of freedom still works. This means the quantum picture of the torsional transition for protein and RNA folding remains to be effective and the quantum calculation based on this picture is reasonable.

Eqs (7)–(10) are basic equations for conformational transition. To obtain quantitative result one should calculate the number of torsion modes *N* in advance. *N* describes the coherence degree of multi-torsion transition in the folding. For two-state protein folding we assume that *N* can be obtained by numeration of all main-chain and side-chain dihedral angles on the polypeptide chain except those residues on its tail which does not belong to any contact. A contact is defined by a pair of residues at least four residues apart in their primary sequence and with their spatial distance no greater than 0.65 nm. Each residue in such contact fragment contributes 2 main-chain dihedral angles and, for non-alanine and -glycine, it contributes 1 - 4 additional side-chain dihedral angles [19] (Table S1 in the Supplementary material). For RNA folding, we assume the quantum transition occurs between compact (yet disordered) intermediate and folding state [20] or between primary and secondary structures of the molecule [21]. The torsion number can be estimated by chain length. Following IUB/IUPAC there are 7 torsion angles for each nucleotide, namely

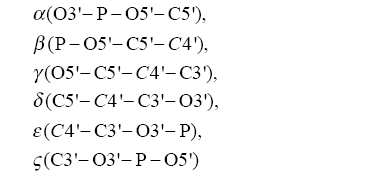

and

*χ* (O4′- C1′- N1- C2) (for Pyrimidine) or *χ*(O4′- C1′- N9 - C2) (for Purine), of which many have more than one advantageous conformations (potential minima). If each nucleotide has *q* torsion angles with multi-minima in potential then the torsion number *N*=*qL*, where *L* is chain length of RNA.

## 3 Results and Discussions

### 3.1 Law on the temperature dependence of folding rate

#### 3.1.1 Deduction of the temperature dependence law

By using the relation of Δ*G* with Δ*E* (Eq (9)) and by the expansion of Δ*E* at melting temperature *T_c_*, Δ*E*(*T*)=Δ*E*(*T_c_*)+*m*(*T* -*T_c_*) we obtain a linear relation between free energy change Δ*G* and temperature. This linearity can be tested rigorously by experiments (Fig S1 in the Supplementary material). Set 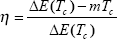. Assuming that 1) the measured value of folding free energy decrease is denoted by Δ*G_f_* and 2) the measurement is carried out at temperature *T_f_*, then one has [13][14]

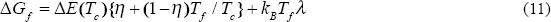

Inserting above equations into (9) and using Eqs (7) and (8) we obtain the temperature dependence of logarithm rate

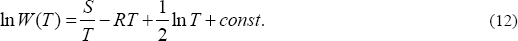

where *const* means temperature-independent term and

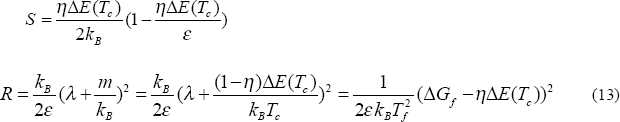

with 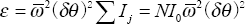. Finally we have

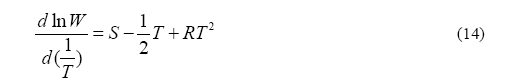

It means the non-Arrhenius behavior of the rate–temperature relationships. The relation was tested for 16 two-state proteins whose temperature dependence data were available. The statistical analyses were made in [13] [14].

Figure 1 gives two examples. The comprehensive comparisons of the theoretical prediction with experimental folding/unfolding rate versus temperature can be found in [13]. The strong curvature of folding rate on Arrhenius plot is due to the *R* term in Eq (12) which comes from the square free energy (Δ*G*)^2^ in ln*W*. The good agreement between theory and experiments affords support to the concept of quantum folding. Moreover, in this model the universal non-Arrhenius characteristics of folding rate are described by only two slope parameters *S* and *R* and these parameters are related to the known folding dynamics. All parameters related to torsion potential defined in this theory (such as torsion frequency 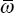 and 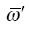, averaged angular shift *δθ* and energy gap Δ*E* between initial and final torsion potential minima, etc) can be determined, calculated consistently with each other for all studied proteins [13]. Furthermore, in this theory the folding and unfolding rates are correlated with each other, needless of introducing any further assumption [6].

**Figure 1.**
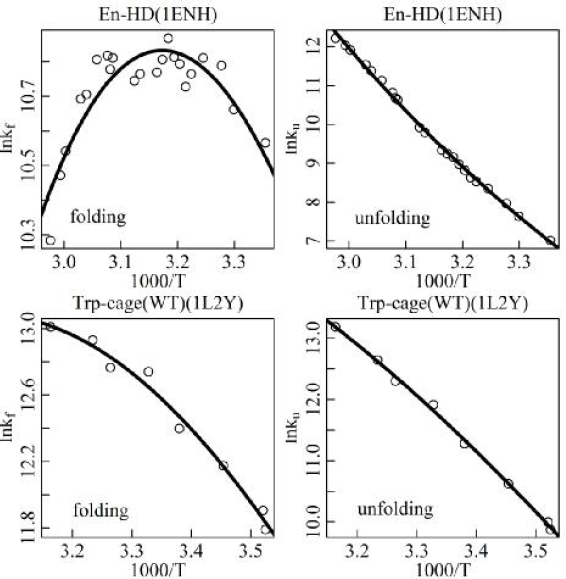
Model fits to overall folding rate *k_f_* and unfolding rate *k_u_ vs* temperature 1000/*T* for protein En-HD (PDB code 1ENH) and Trp-cage(WT) (PDB code 1L2Y). Experimental logarithm folding rates are shown by “o”, and solid lines are theoretical model fits to the folding rate (*k_f_* in unit s-^1^, *T* in unit Kelvin). Experimental rates are taken from [22, 23].

#### 3.1.2 Temperature dependence of free energy and relations among folding parameters

The temperature dependence of free energy change Δ*G* plays an important role in the deduction of temperature dependence law for protein folding. We shall make deeper analysis on this point and, based on this analysis, deduce more relations among folding and unfolding parameters on Arrhenius plot. Assuming the free energy changes Δ*G* in a temperature interval lower than *T_c_* have been measured and expressed as

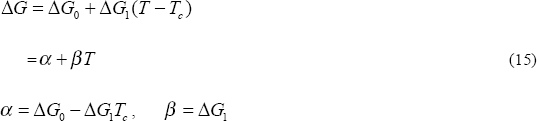

The linear relations are tested by experiments for many proteins (Figure S1). By using *α* and *β* given from experiments we can re-deduce the temperature dependence of folding rate. In fact, by inserting Eq (15) into Eq (7) one easily obtains Eq (12) and in the equation the slope and curvature parameters are given by

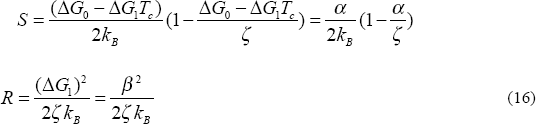

Eliminating *ζ* in Eq (16) a universal relation between *R* and *S* is deduced

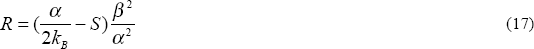

The temperature dependence of unfolding rate *W_u_* is easily obtained by the replacement of Δ*G* by -Δ*G* and 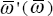 by 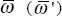 in folding rate *W*. As Eq (12) for folding rate the unfolding rate can be expressed as

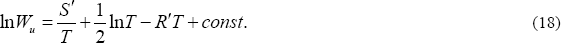

Following the similar deduction of Eq (16) and (17) the slope and the curvature parameters of unfolding rate are obtained,

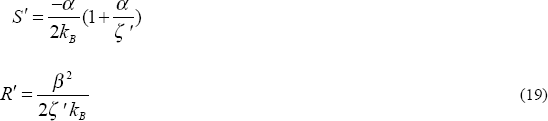

and, by eliminating *ζ*

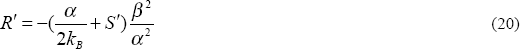

With the aid of known *α* and *β* from the free energy temperature dependence Eqs (17) and (20) give constraints on slope and curvature parameters for folding/unfolding rate. This provides a new checkpoint for the present quantum folding theory.

In addition to Eqs (17) and (20), by use of the temperature dependence of free energy, Eq (15), one can deduce relationship between folding and unfolding rates. In fact, Eqs (7)–(10) can be used for unfolding as well as for folding. By the replacement of Δ*G* by -Δ*G* and 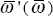 by 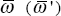 in *W* we obtain unfolding rate *W_u_*. We have

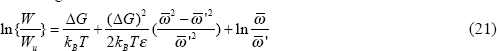

Eq (21) means the condition of dynamical balance *W* =*W_u_* for protein folding is slightly different from the usual equilibrium condition Δ*G*=0 for chemical reaction due to the unequal bias samplings in frequency space, 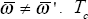 is defined by *W* =*W_u_*. Eq (21) means *T_c_* and Δ*G*_0_ satisfying

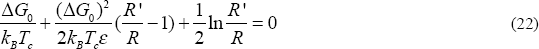

In deducing above equation 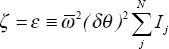 in (16) and 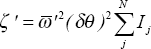 in (19) have been used. As Δ*G*_0_ has been known the ratio *R*′/*R* can be solved from Eq (22). The result can be compared with the experimental *R* and *R*′ obtained from quadratic fits to experimental rates of folding and unfolding rates. This provides another checkpoint for the quantum folding theory.

The free energy parameters *α,β*, Δ*G*_0_, and the calculating results from Eqs (17) and (22) are listed in Table 1.

**Table 1.** Temperature-dependence parameters about free energy and folding rate for 15 proteins

| PDB code | $\alpha(10^{-20} \text{ J})$ | $\beta(10^{-20} \text{ J/K})$ | $\Delta G_0/k_B T_c$ | $\varepsilon/k_B T_c$ | $\log(R'/R)_{\text{pred}}$ | $\log(R'/R)$ | $R_{\text{pred}}/R$ |
| --- | --- | --- | --- | --- | --- | --- | --- |
| 1bdd | 20.10 | -0.05579 | 0.03140 | 9.30 | -0.027 | -0.025 | 1.01 |
| 1div<sub>n</sub> | 15.77 | -0.04299 | 0.94765 | 4.63 | -0.75 | -0.019 | 0.96 |
| 1e0l | 17.05 | -0.05216 | -0.01401 | 12.67 | 0.012 | -0.034 | 1.19 |
| 1enh | 20.79 | -0.06406 | -0.06577 | 10.46 | 0.057 | -0.016 | 1.16 |
| 1l2y(p12w) | 10.54 | -0.03196 | -0.08485 | 4.00 | 0.073 | -0.038 | 1.04 |
| 1l2y(wt) | 7.927 | -0.02526 | -0.00840 | 2.75 | 0.009 | -0.013 | 1.22 |
| 1lmb(wt) | 33.50 | -0.1002 | 0.07203 | 9.73 | -0.062 | -0.019 | 1.12 |
| 1lmb(g46a) | 37.87 | -0.1114 | -0.39823 | 13.22 | 0.215 | -0.026 | 1.07 |
| 1lmb(sa37g) | 29.42 | -0.08996 | -0.19194 | 6.06 | 0.166 | -0.020 | 1.07 |
| 1pin(wt) | 24.16 | -0.07282 | -0.03545 | 6.28 | 0.031 | -0.048 | 1.05 |
| 1pin(s18g) | 23.48 | -0.07139 | -0.17282 | 5.33 | 0.149 | -0.048 | 1.05 |
| 1pin(n26d) | 18.88 | -0.06075 | -0.03087 | 9.91 | 0.027 | -0.025 | 1.17 |
| 1prb | 27.73 | -0.07431 | 0.02403 | 9.71 | -0.021 | -0.034 | 1.06 |
| 2a3d | 21.15 | -0.06110 | -0.10796 | 14.84 | 0.094 | -0.013 | 1.19 |
| 2pdd | 22.27 | -0.06840 | -0.06313 | 2.26 | 0.055 | -0.043 | 1.02 |

From Tab 1 we find: 1) *R_pred_*/*R* takes a value near 1. The differences between two sides of Eq (17), the relative errors in predicting *R* by use of Eq(17), are in the range of 1% to 22% for different proteins and smaller than 10% for most of them. Thus the constraint equation between slope and curvature parameters on Arrhenius plot are proved.

2) (*R*′/*R*)_pred_ calculated from Eq (22) are compared with *R*′/*R* given from the statistical analyses of folding/unfolding rate data. The difference between log(*R*′/*R*)_pred_ and log(*R*′/*R*) is generally smaller than 0.23 apart from one protein (1div) as seen from Tab 1. This means the ratio of (*R*′/*R*)_pred_ to (*R*′/*R*) is generally smaller than 1.7 for most proteins. Considering the possible errors in the experimental determination of *R*′ and *R* the theoretical prediction based on equation (22) is acceptable. Moreover, (*R*′/*R)_pred_* given in Tab 1 takes a value between 0.86 and 1.64 apart from 1div From 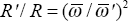 we estimate the frequency ratio 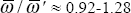

3) Δ*G*(*T_c_*) ≡ Δ*G*_0_ ≠ 0, which is deduced from the linear temperature-dependence of free energy. If Δ*G*_0_ were 0 then *R*′ would equal *R* from Eq (22). So the inequality between *R*′ and *R* indicated by experiments would require A*G*_0_ ≠ 0. Of course, Δ*G*_0_ is a small quantity in the order of 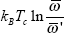. However, in the accurate determination of free energy at a given temperature *T* near *T_c_* one should consider the non-vanishing contribution from Δ*G*_0_ term.

So far we have discussed the temperature dependence of protein folding. The temperature dependence law Eq (12) deduced from quantum folding theory can also be used for RNA folding. For RNA molecule the temperature dependence of folding rate was observed for yeast tRNA^phe^ [24]. They measured the logarithm folding rate ln*k_f_* versus 1/*T* between 28.5°C and 34.8°C. The Arrhenius plot shows a straight line in this temperature interval but large standard deviation existing at low temperature end. From the experiments on protein folding, the strong curve of the ln*k_f_* - 1/*T* relation only occurs in a temperature interval of several tens degrees. We expect more accurate measurements within a large enough temperature interval will be able to exhibit the non-Arrhenius peculiarity of the temperature dependence of the RNA folding rate.

### 3.2 Relation of fast-variable factor of folding rate with respect to torsion number *N*

For a large class of conformational change problems, for example in the conventional protein and RNA folding, the chemical reaction and electronic transition are not involved and the fast variables include only bond lengths and bond angles of the macromolecule. In this case an approximate relation of the fast-variable factor 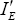 with respect to torsion number *N* can be deduced. When the kinetic energy in 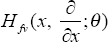 is neglected as compared with interaction potential *U_fv_* one has

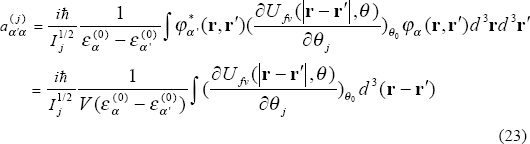

In the above deduction of the second equality the fast-variable wave function *φ_α_*(**r,r′**) has been assumed to be a constant and normalized in the volume *V*. As the energy and volume *V* are dependent of the size of the molecule one may assume energy 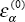 and *U_fv_* proportional to the interacting-pair number (namely *N*^2^) and *V* proportional to *N*. However, because only a small fraction of interacting-pairs correlated to given 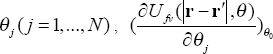 does not increases with *N*. So, one estimates 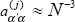. On the other hand, the integral 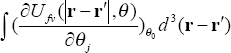 may depend on the molecular structure. For example, the high helix content makes the integral increasing. It was indicated that a protein with abundant *α* helices may have a quite oblong or oblate ellipsoid, instead of spheroid, shape and this protein has higher folding rate[12][13]. Therefore, apart from the factor *N*^−3^ there is another structure-related factor in 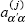 though the latter is *N*-independent. Assuming *M* proportional to *N*, one obtains

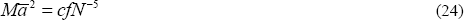

where *f* is a structure-related shape parameter. It means the fast-variable factor 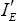; is inversely proportional to *N*^5^. With 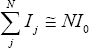 and Eq (24) inserted into Eq (7)–(8) we obtain the relation of logarithm rate with respect to *N* and Δ*G* [25]

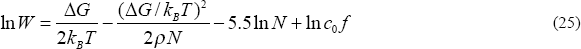

where 

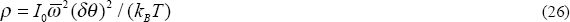

is a torsion-energy-related parameter and 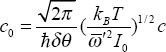 is an *N*-independent constant.

The relationship of In*W* with *N* given by Eq (25) can be tested by the statistical analyses of 65 two-state protein folding rates *k_f_*. We found that the theoretical logarithm rate In*W* is in good agreement with the experimental ln*k_f_* (Fig 2). The figure is plotted for *ρ*=0.097 and it gives the correlation between theoretical and experimental rates *R*=0.7818 and slop of regression line 1.109. For any *ρ* between 0.06 and 0.1 the basically same results are obtained, for example, *R*=0.7537 and slop=1.044 for *ρ* =0.069, *R*=0.7396 and slop=0.997 for *ρ* =0.06.

**Figure 2.**
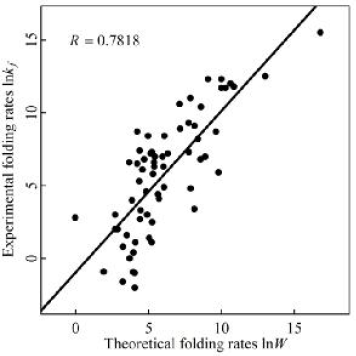
Comparison of theoretical folding rates In*W* with experimental folding rates in*k_f_* for 65 proteins. The experimental rates *k_f_* are taken from Table 1 of ref [13]. The straight line is the linear regression between In*W* and In*k_f_*. In calculation of theoretical In*W* by use of Eq (25) the shape parameter *f* is taken as follows: *f*=81 for (*L_α_* - *L_β_*)/*L* ≥ 0.6, *f*=25 for 0.3≤(*L_α_* - *L_β_*)/*L*< 0.6, and *f*=1 for (*L_α_* - *L_β_*)/*L*< 0.3.( *L_α_* and *L_β_* are the number of residues in *α* helix and *β* sheet, respectively, and *L* is the total number of folded residues). Our experience shows that the different choice of *f*-value in the intermediate region is insensitive to the statistical result.

### 3.3 Relation of free energy Δ*G* with respect to torsion number *N*

To find the relation between free energy Δ*G* and torsion number *N* we consider the statistical relation of free energy combination 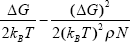 that occurs in rate equation (7) or (25). Set

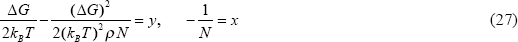

The linear regression between *y* and *x* is given by *y* =*A* + *Bx* where *A* and *B* are two statistical parameters describing free energy distribution in the dataset. We will test the linear relation in protein folding dataset [12] [13]. Due to the ignorance of the accurate ρ-value for each protein one can test the relation by using the single-*ρ*-fit (assuming a single *ρ*-value to deduce a linear regression) at first, then compare the fitting results and find the best-fit *ρ*-value and the corresponding free-energy statistical parameter *A* and *B*. The statistical results (correlation *R* between *y* and *x* and parameter *A* and *B*) in two-state protein dataset are listed in Table 2 From Table 2 we find the correlation *R* between *y* and *x* is near to 0.8 for *ρ*=0.065∼0.075 and reaches maximum at *ρ*=0.069 where *R*=0.7966. Thus, by single-*ρ*-fit we obtain the best-fit statistical relation of free energy for two-state proteins as

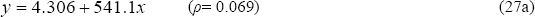

**Table 2.** Free energy parameters determined by linear regression

| $\rho$ | 0.05 | 0.060 | 0.065 | 0.069 | 0.075 | 0.097 |
| --- | --- | --- | --- | --- | --- | --- |
| $R$ | 0.6573 | 0.7759 | 0.7934 | 0.7966 | 0.7919 | 0.7507 |
| $A$ | 3.194 | 3.867 | 4.126 | 4.306 | 4.540 | 5.152 |
| $B$ | 428.0 | 496.4 | 522.8 | 541.1 | 564.9 | 626.9 |

$A$ and $B$ - free energy parameter, $R$ - correlation coefficient. $R$ reaches maximum at $\rho=0.069$ .

(Figure 3a). In above discussion the single-*ρ*-fit has been used. Evidently, as the variation of *ρ* for different proteins is taken into account the linear regression between free energy combination *y* and torsion number *x* will be further improved.

**Figure 3.**
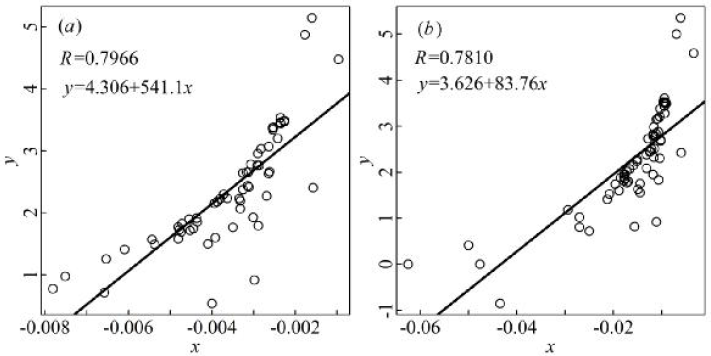
Statistical relation of free energy for two-state proteins. Experimental data are taken from 65- protein set [12][13]. Five proteins in the set denatured by temperature have been omitted in our statistics. In (*a*) *y* and *x* are defined as Eq (27); in (*b*) *y* and *x* defined as Eq (28).

Because *N* increases linearly with the length *L* of polypeptide chain, instead of Eq (27), by setting

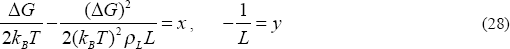

we obtain the best-fit statistical relation of free energy for two-state proteins (Fig 3b) as

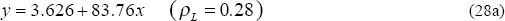

and the correlation coefficient *R*=0.781.

About the relationship of free energy Δ*G* with torsion number *N* or chain length *L* several proposals were proposed in literatures. One statistics was done by assuming the linear relation between Δ*G* 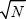 and 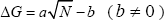 [13]. Another was based on the assumed relation of Δ*G vs* (*Lg* + *σB_L_L*^2/3^) [12]. By the statistics on 65 two-state proteins in the same dataset we demonstrated the correlation *R* between free energy and *N* or *L* is 0.67 for the former and 0.69 for the latter [13], both lower than the correlation shown in Fig 3. On the other hand, the proposal of free energy scaled as 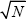 or 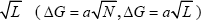 [17] seems not agree with protein experiments. The free energy combination of 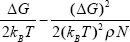 occurring in the folding rate is an important peculiarity of the present theory. We shall use the statistical relation of the free-energy combination versus *N* in the following studies on RNA folding.

### 3.4 Relation of folding rate with respect to torsion number *N* and its test in RNA folding dataset

In virtue of Eqs (25) (27) we obtain an approximate expression for transitional rate In*W* versus *N* for protein folding

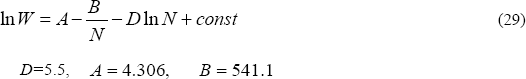

Here *const* is an *N*-independent constant but dependent of molecular shape (through the factor *f* contained in it). Neglecting the shape-dependence and putting *N* proportional to chain length *L* we re-write the relation as

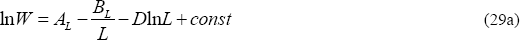

and compare it with 65- protein dataset the correlation between In*W* and experimental ln*k_f_* is *R*=0.722 as *A_L_*=3.626, *B_L_*=83.76 (Eq 28a) and *D*=5.5 are taken.

The quantum folding theory of protein is applicable in principle for each step of the conformational transition of RNA molecule. Although recent experiments have revealed the multi-stages in RNA collapse, the final search for the native structure within compact intermediates seems a common step in the folding process. Moreover it exhibits strong cooperativity of helix assembly [15][20]. Because the collapse transition prior to the formation of intermediate is a fast process and the time needed for the former is generally shorter than the latter [20], the calculation of the transition from intermediate to native fold can be directly compared with the experimental data of total rate. Furthermore, for RNA folding the *const* term in Eq (29) can be looked as a real constant if the variation of structure-related shape parameter *f* is neglected in the considered dataset. By using *N*=*qL* (*L* is the chain length of RNA) we have

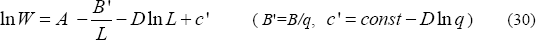

Eq (30) is deduced from quantum folding theory with some statistical consideration and it predicts the relation of folding rate versus chain length: the rate *W* increasing with *L*, attaining the maximum at *L_max_*=*B*′/*D*, then decreasing with power law *L^−D^*.

In a recent work Hyeon and Thirumalai [16] indicated that the chain length determines the folding rates of RNA. They obtained a good statistical relation between folding rates and chain length *L* in a dataset of 27 RNA sequences. Their best-fit result is

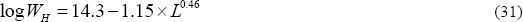

Both equations (30) and (31) give the relation between RNA folding rate and chain length. Comparing the theoretical folding rates In*W* or In*W_H_* with the experimental folding rates ln *k_f_* in 27 RNA dataset the results are shown in Figure 4. We found Eq (30) can fit the experimental data on RNA folding rate equally well as Eq (31). By using the best-fit value of *B* ’ and *D* the correlation between ln *W* (calculated from Eq 30) and ln *k_f_* is *R*=0.9729 (Fig 4a), while the correlation between In*W_H_* (calculated from Eq 31) and ln *k_f_* is *R*=0.9752(Fig 4b). However, in Fig 4b the slope of the regression line is 1.03 and the line deviates from origin by -0.36, while in Fig 4a the slope is 1.0001, very close to 1 and the line deviates from origin only by -0.0012. The reason lies in: although two equations have the same overall accuracy in fitting experimental data, but for large *L* cases the errors *Er* =|log*W*-log*k_f_*| calculated from Eq (30) are explicitly lower than *Er_H_* =|log*W_H_*-log*k_f_*| from Eq (31) (Table 3). It means the folding rate lowers down with increasing *L* as *L^−D^* (*D* ≅ 5.5) at large *L* (a long-tail existing in the *W-L* curve) rather than a short tail as 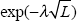 assumed in [16]. The long-tail form of folding rate can be used to explain some small- probability events in pluripotency conversion of gene [25].

**Table 3.** Errors of RNA folding rates in two theoretical models compared with experimental data

|  | 1 | 2 | 3 | 4 | 5 | 6 | 7 | 8 |
| --- | --- | --- | --- | --- | --- | --- | --- | --- |
| $L$ | 125 | 160 | 205 | 225 | 368 | 377 | 409 | 414 |
| $k_f(\text{s}^{-1})$ | 6 | 2 | 10.5 | 6.5 | 0.03 | 0.011 | 0.008 | 0.013 |
| $Er_H$ | 0 | 0.17 | 3.27 | 3.37 | 1.52 | 0.71 | 1.06 | 1.66 |
| $Er$ | 0 | 0.18 | 3.15 | 3.17 | 0.44 | 0.42 | 0.30 | 0.26 |

1, hairpin ribozyme [26][27]; 2; P4-P6 domain (*Tetrahymena* ribozyme)[28]; 3, *Azoarcus* ribozyme [27][29]; 4, *B. subtilis* RNase P RNA catalytic domain[30]; 5, Ca.L-11 ribozyme[31]; 6, *E. coli* RNase P RNA[32]; 7, *B. subtilis* RNase P RNA[32] and 8, *Tetrahymena* ribozyme[27][33]. Experimental rates $k_f$ 's can be found in literatures [26-33]. The errors of eight RNAs with lengths larger than 120 in 27-RNA dataset are analyzed in the table. The errors of the first RNA hairpin ribozyme are normalized to zero in two models.

**Figure 4.**
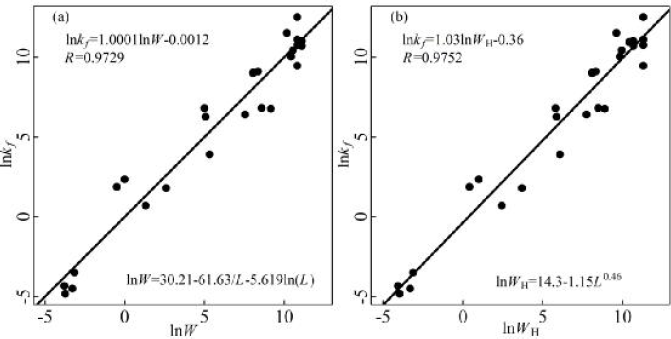
Comparison of experimental folding rates In*k_f_* with theoretical folding rates ln*W* (Fig 4a) or ln*W_H_* (Fig 4b) for 27 RNA molecules. Experimental rates are taken from Table 1 in ref [16]. Theoretical rates are calculated from Eq (30) and (31) respectively.

There are two independent parameters in RNA folding rate Eq (30), *B*′ and *D*, apart from the additive constant. As seen from Fig 4a we obtain the best-fit *D* value *D_f_*=5.619 on the 27-RNA dataset, close to *D*=5.5 predicted from a general theory of quantum folding. Simultaneously we obtain the best-fit *B*′ value 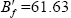. The 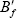 value derived from RNA folding can be compared with the *B_L_*=83.76 from protein folding (Eq (29a)). Notice that *B_L_* or *B*′ represents the contribution from free energy square term in logarithm folding rate. The RNA folding free energy is typically 2 to 4 kcal/mol [20,34,35] while the folding free energy for most proteins in 65-protein dataset is between 1 and 4.6 kcal /mol. (Table 1 in ref [13]). The two free energy values are near each other. It explains 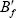 near *B_L_*.

The 27-RNA dataset [16] contains data of experimental folding rates measured in different processes. The dataset is inhomogeneous, including several subsets, one subset of total folding rates for some RNAs and another subset of rates of secondary structure formation for other RNAs, etc. One wondered why the simple formula such as Eq (30) can fit the experiments so well. The reason may be in: all RNA folding processes can be looked as the quantum transition between conformational states and this common mechanism has been quantitatively described by a unifying equation; furthermore, all *D*’s occurring in the equation for different subsets are near to 5.5 by theoretical grounds. So one can use Eq (30) with single parameters *B*′, *D* and an additive constant to fit the experimental folding rates. However, the dissimilarity of free energy parameters *B* (and *A*) and the variation of structure-related parameter *f* in different subsets do exist. The differences of these parameters have been neglected in the simplified equation (30). A more rigorous comparison between theory and experiments should take these differences into account. For example, as the nucleotide G in the tetraloop hairpin UUCG is substituted by 8-bromoguanosine 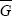, the folding rate of gcUUC 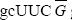 gc is 4.1-fold faster than gcUUCGgc [21]. Both samples were collected in the dataset [16]. They have the same chain length *L*=8 but different rate *k_f_*. This cannot be explained by Eq (30). However, if the variations of the free energy change Δ*G* and the structure-related parameter *f* are incorporated in a more rigorous equation (as in the original Eq (25)) then the detailed difference in the folding rate for these two samples will be interpreted in a natural way.

#### Remarks

We have studied the folding of two-state proteins and RNA molecules from the point of quantum transition. Dislike two-state protein the RNA folding is a multi-stage process. Both the folding from compact intermediate to native fold and the folding of secondary structure formation can be studied from quantum folding theory. However, to study the collapse transition as a whole for RNA molecule needs further development of the quantum model. The problem can be compared with the multi-state protein folding. For multi-state protein one may assume the folding is a mutual process of several quantum transitions in different domains and that some time delays exist between these transitions [36]. The similar idea might be introduced in the study of the total folding rate of the RNA molecule.

To test quantum folding theory we suggest make experimental study directly on protein and RNA photo-folding. The stimulated photo-folding rates and the resonance fluorescence cross section have been calculated for protein in [37]. These results can be generalized to RNA. The particular form of the folding rate–temperature relation and the abundant spectral structure in protein and RNA photo-folding will afford direct evidence on the existence of a set of quantum oscillators of low frequency and the quantum transitions between molecular conformations.

## 4 Summary of Quantitative Relations

Based on quantum theory of conformation change of biomolecule the following quantitative results are deduced:

1) A law on the temperature dependence of folding rate

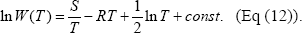 The rate formula successfully explained the non-Arrhenius peculiarity of protein folding in a natural way.
2) The relation between slope *S* (*S*′) and curvature *R*(*R*′) on Arrhenius plot for protein folding/unfolding

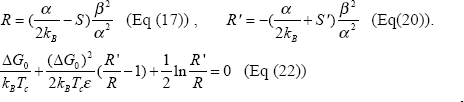
3) The linear relation *y*=*Ax*+*B* between free energy 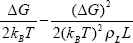 (*y*) and chain length -1/*L* (*x*) (Eq (28) and (28a)).
4) A statistical formula on protein and RNA folding rate dependent of torsion number (chain length)

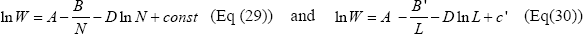 The theoretical predictions are in accordance with the experimental data on two-state-protein and RNA folding The partial success of the present study on protein and RNA folding from a simple unified theory reveals the existence of a common quantum mechanism in the conformational transition of biomolecules.

## Author Contributions

L.L proposed the theoretical model and deduced the folding formula. J.L. made the statistical analyses on protein and RNA folding.

## Acknowledgment

The work was supported by the Inner Mongolia Autonomous Region Natural Science Foundation (No. 2015MS0331). The author would like to thank Drs Zhao Judong and Zhang Ying for numerous discussions on protein data analyses, and Dr Bao Yulai for his help in literature searching.

## Supplementary Material

**Table S1.** The number of side-chain dihedral angles *n*′ for 20 amino acids

| amino acids | Ala | Arg | Asn | Asp | Cys | Gln | Glu | Gly | His | Ile |
| --- | --- | --- | --- | --- | --- | --- | --- | --- | --- | --- |
| $n'$ | 0 | 4 | 2 | 2 | 1 | 3 | 3 | 0 | 2 | 2 |
| amino acids | Leu | Lys | Met | Phe | Pro | Ser | Thr | Trp | Tyr | Val |
| $n'$ | 2 | 4 | 3 | 2 | 2 | 1 | 1 | 2 | 2 | 1 |

**Figure S1.**
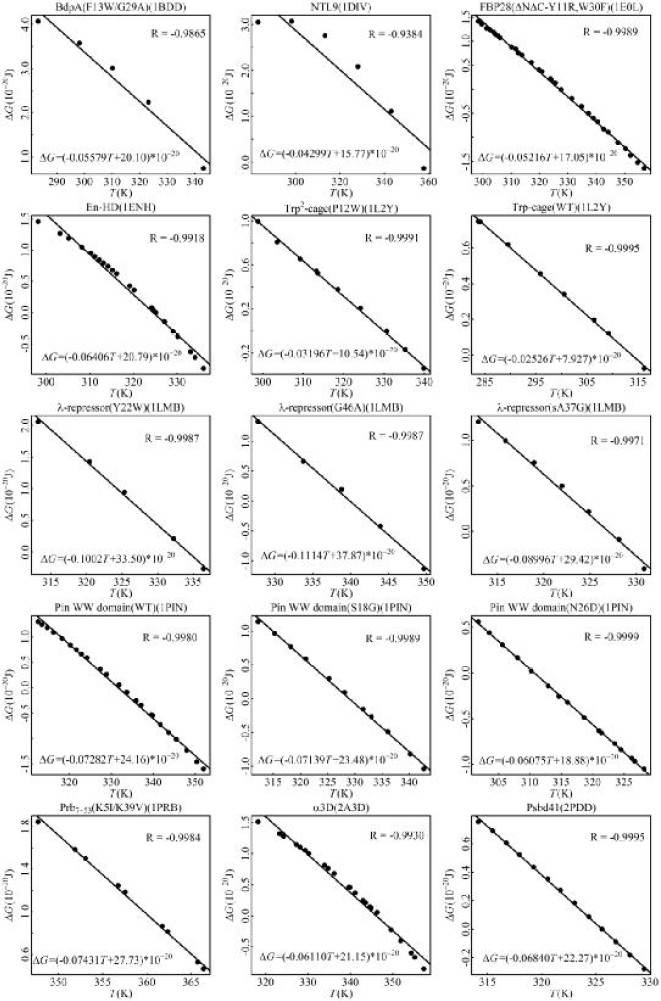
Linear relation between protein free energy Δ*G* and temperature. Experimental free energies Δ*G* changing with temperature are plotted for 15 proteins. The protein 1iet in 16 proteins is not shown in this figure due to the scarcity of data. The regression analysis shows a good linear relation existing between Δ*G* and *T* for each protein. Δ*G* in unit J, *T* in unit Kelvin. References on the experimental data can be found in Tab 2 of ref [13].

